# Energetic Evolution of Cellular Transportomes

**DOI:** 10.1101/218396

**Authors:** Behrooz Darbani, Douglas B. Kell, Irina Borodina

## Abstract

Transporter proteins mediate the translocation of substances across the membranes of living cells. We performed a genome-wide analysis of the compositional reshaping of cellular transporters (the transportome) across the kingdoms of bacteria, archaea, and eukarya. We show that the transportomes of eukaryotes evolved strongly towards a higher energetic efficiency, as ATP-dependent transporters diminished and secondary transporters and ion channels proliferated. This change has likely been important in the development of tissues performing energetically costly cellular functions. The transportome analysis also indicated seven bacterial species, including *Neorickettsia risticii* and *Neorickettsia sennetsu*, as likely origins of the mitochondrion in eukaryotes, due to the restricted presence therein of clear homologues of modern mitochondrial solute carriers.

## Introduction

The expansion of life on Earth has involved competition among organisms for limited energy resources [1]. Arguments have been made in favour of growth rate over growth efficiency in organisms competing within a specific niche [2], which implies a natural selection towards an improved ability to capture and utilize the available energy sources for survival and reproduction [3,4]. Darwinian evolutionary theory originally covered only phenotypic improvements at the organismal level, but we nowadays also recognize the importance of molecular and cellular events such as the acquisition of mitochondria by eukaryotes. This enabled an increase of eukaryotic cell size and complexity due to a more “efficient” generation of the cellular fuel ATP [5,6]. Cells need to allocate considerable resources to energize their transportomes. For example, brain neurons account for approximately 20% of the basal metabolic rate in humans, mostly for the movement of ions across neuronal membranes [7]. In general, a metabolic cost of up to 60% of the total ATP requirement in organisms is estimated for the activity of their transportomes [8,9]. Thus, it would be reasonable to hypothesize that an improved energetic performance of the transportome might be selected for over the course of evolution.

Despite the importance of cellular transportomes (also reported as the second largest component of the human membrane proteome [10]), transporters are surprisingly understudied [11]. Additionally, the presence of substrate-binding proteins as the partners of a subset of membrane transporters [12] makes the cellular transport machinery more complicated than if we consider only the transporters themselves. The tightening of porous and leaky primordial envelopes such that they did not let in (and could learn to efflux) all kinds of substances [13,14] has been proposed as a turning point for membrane transporters to co-evolve together with lipid bilayer membranes [15]. Different classes of transporters, each including several transporter superfamilies, share a common ancestral family of peptides which carry 1-4 transmembrane domains and form homo- and hetero-oligomer channels [16–19]. Intragenic duplication and triplication have been the major events promoting the diversification of transporter proteins [17,19]. Classical evolutionary theory on the basis of natural selection proposed by Charles Darwin [3,4] explains how the random variability of the genome as the diversification force has given the chance for specialisation, improved performance, and adaptation to the continuously changing biosphere with limited energy resources. Here, we therefore annotated the cellular transportomes of bacteria, archaea and eukarya and analysed their composition with a focus on the energetic efficiency. The analyses include all the three classes of transporters, *i.e.,* ion channels, secondary transporters, and ATP-dependent transporters. To translocate substrates, ATP-dependent transporters bind and hydrolyse ATP [20], ion channels form pores for selective diffusion of ions and small molecules, and the secondary transporters shuttle substrate molecules across biological membranes either through energy independent facilitated diffusion or via exploiting membrane electrochemical potential gradients through uniport, symport and antiport [21].

## Results and Discussion

To study the compositional changes of transportomes, we analysed the transportomes of 249 evolutionarily distant species (of which 222 were annotated in this study) from archaea, bacteria and eukarya. These included 126 prokaryotic species (from 16 taxonomic phyla and 60 taxonomic orders), 30 primitive eukaryotes (different species from Alveolata, Kinetoplastida and Amoebozoa), 30 algal and plant species, 30 fungal species, and 33 animal species (See S1 Data). The transportomes were annotated using the Transporter Automatic Annotation Pipeline at TransportDB [22]. Notably, the species had large differences in the size of both their genomes and their transportomes (Fig 1a, b and S1 Data).

**Fig 1.**
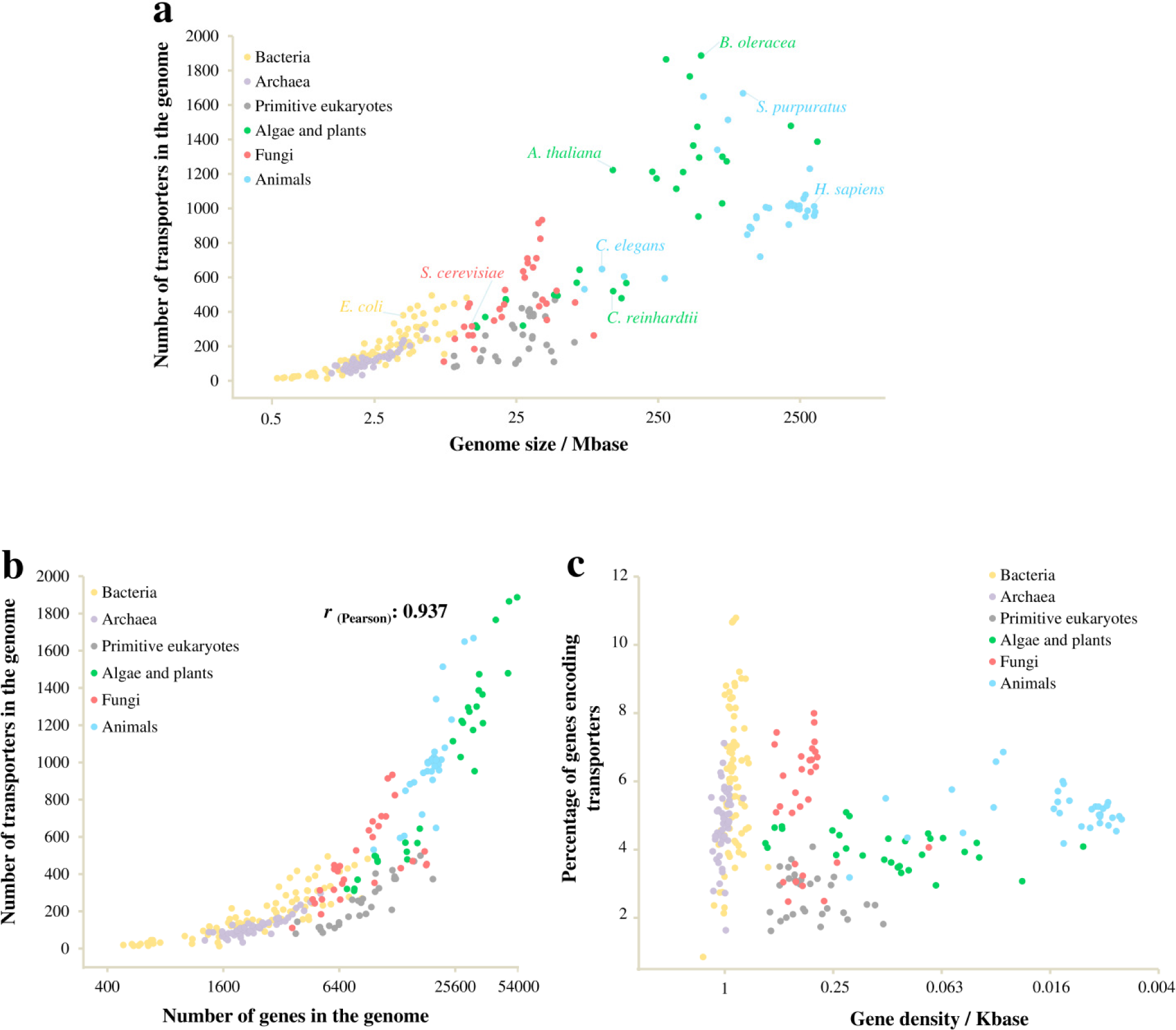
Transportomes differ in size among species and evolutionarily distant domains of life. **(a)** The number of membrane transporters per organism in relation to the genome size. **(b)** The number of membrane transporters per genome in relation to the number of total genes. **(c)** Percentage of transporter-coding genes in relation to gene density.

We found that the sizes of an organism’s transportome tends to increase with its genome size, though there are considerable intra-kingdom variations (Fig 1a) (see also [23]). In particular, primitive eukaryotes have small transportomes with 100-500 members, relative to their genome sizes of tens of Mb when compared to the bacteria with modestly sized genomes (less than 10 Mb). Ion channels, secondary transporters, and primary active transporters were found in each of the analysed domains of life (See S1 Data). This indicates a very early appearance for these three classes of transporters, possibly in a common ancestor.

In agreement with previous reports [24,25], the genome size had a higher rate of enlargement than did the gene number, resulting in a decreased gene density over the course of evolution (Fig 1c). The transportome enlargement was found to be well correlated with the increase in the gene number (*r*_(Pearson)_ = 0.937) with the exception of primitive eukarya, where the transportome-encoding proportion of genes was the lowest (Fig 1b, c). Of most interest, the composition of the transportomes changed from prokaryotes to eukaryotes and also among the eukaryotic kingdoms along with the transportome enlargement (Fig 2a). Specifically, we found higher intra-genomic frequencies (frequency relative to the total number of genes in the genome) of ATP-dependent transport classes in prokaryotes than eukaryotes. An opposing trend was found for low-energy-demanding transport classes. This indicates different rates of gene proliferation among the evolved transporter classes; low-energy-demanding transporter families have expanded at a higher rate. Notably, the transportomes of primitive eukaryotes also represent a transition state between prokaryotic and higher eukaryotic kingdoms (Fig 2a).

**Fig 2.**
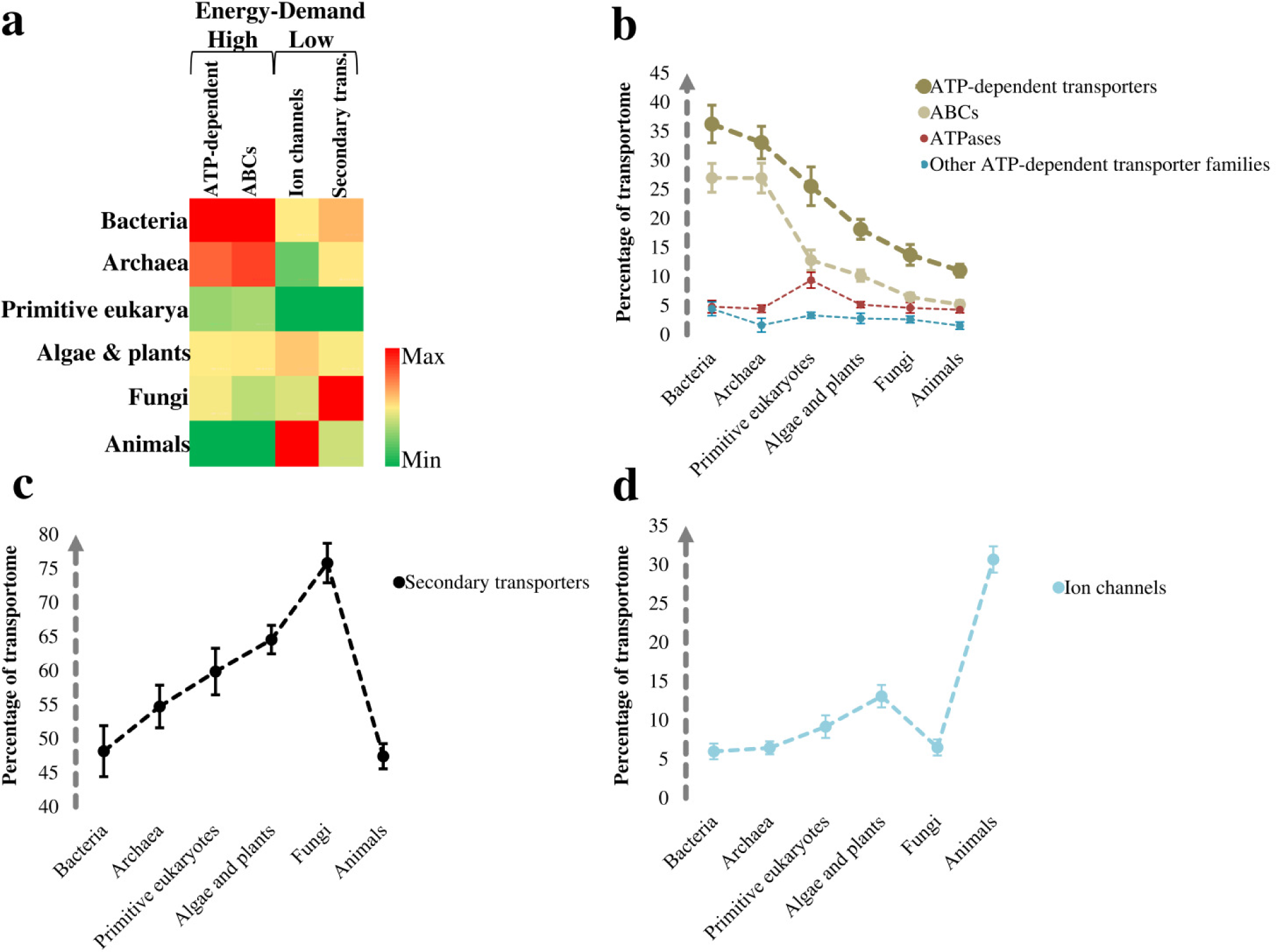
The evolutionary dynamics of transportomes composition. **(a)** Heat-map representation of the changes in the number of members of the transporter classes. To calculate the intra-genomic frequencies, the numbers of transporter members are normalized to the total number of genes per genome. The heat-map is drawn for each class of ion channels, secondary transporters, and ATP-dependent transporters and therefore colors are not comparable between the classes. **(b)** The fraction of ATP-dependent transporters in the transportomes. All of the variations of ATP-dependent transporters and ABC superfamily except the difference between bacteria and archaea are significant at 0.1% level. **(c)** The fraction of secondary transporters in the transportomes. Only the difference between bacteria and animals is not statistically significant (*p* = 0.635). **(d)** The fraction of ion channels in the transportomes. All differences, except among fungi, bacteria and archaea, are significant with a *p*-value less than 0.001. The values on panels b-d are shown as mean +/-t-test based 99% confidence intervals. The variations were also confirmed on the arc sin √*x* transformed data (See S1 Data). The family names of the transporters can also be found in the S1 Data.

We further compared the prevalence of secondary transporters, ion channels, and ATP-dependent transporters within the transportomes of each species, and averaged these over the larger taxonomic groups (Fig 2b-d). In general, we observed compositional changes that indicate a positive adaptive selection of the no- to low-cost-flux equilibrative ion channels and carriers, and negative selection of the energetically more expensive ABC transporters. More specifically, we found significant variations in each of the transporter classes (Fig 2b-d). While ca. 27% of all the bacterial and archaeal transporters are ABC transporters, this fraction decreases to 13% in primitive eukaryotes, 10% in algae and plants and a mere 5-6% in fungi and animals (Fig 2b). On the other hand, an increased contribution to the cellular transportome was found for secondary transporters in eukaryotes, particularly in fungi (Fig 2c). Furthermore, animals with ≈ 30% and the green domain of life including algae and plants with ≈ 13% showed higher percentages of ion channels when compared to the bacteria and archaea with only 6%-7% (Fig 2d). During evolution, and in parallel with the genome enlargement and gene pool expansion, each of the three classes of transporters had a chance to contribute proportionally to the expansion of transportomes. By contrast, our results show a preference for the low-energy demanding transporters (ion channels and carriers) over the energy-costly transporter classes (ATP-dependent families, and ABCs in particular) in the transition from prokaryotes to eukaryotes. We defined the energy usage efficiency of a transportome (EUE) as the average required energy per single substrate translocation. We calculated the EUE values for the transportomes studied (more details in the Methods section). The EUE describes the overall energetic performance of transportomes at organismal level and most importantly it does not indicate the total energy consumption by the cellular transportome, because the latter depends also on the flux through individual transporters that is largely unknown. In contrast to the total energy requirements, the EUE is therefore not subjected to spatiotemporal variations.

By comparing the average EUEs of the transportomes across the different domains of life, we found that the EUE has improved in eukaryotes by reductions of up to 0.50 ATP in the average ATP consumption per single transport event mediated by transporters (Table 1).

**Table 1.** Improvement in the energy-usage efficiency (∆EUE) calculated as changes in the average ATP-usage per single transport cycle.

| Domains of life | Bacteria | Archaea | Primitive eukaryotes | Algae and plants | Fungi | Animals |
| --- | --- | --- | --- | --- | --- | --- |
| Bacteria | 0 |  |  |  |  |  |
| Archaea | -0.03 | 0 |  |  |  |  |
| Primitive eukaryotes | -0.16 | -0.14 | 0 |  |  |  |
| Algae and plants | -0.27 | -0.25 | -0.11 | 0 |  |  |
| Fungi | -0.30 | -0.27 | -0.13 | -0.02 | 0 |  |
| Animals | -0.49 | -0.46 | -0.32 | -0.21 | -0.19 | 0 |

The changes ( $\Delta$ EUE) are calculated as $\text{ATP-usage}_{\text{domain of life in the matrix-row}} - \text{ATP-usage}_{\text{domain of life in the matrix-column}}$ (see methods). Negative changes represent the reduction in ATP-usage and improved EUE of transportomes.

Furthermore, the data suggest that animals have mainly relied on the diversification of ion channels, fungi on secondary transporters, and finally, algae and plants on both transporter classes for the evolution of their transportomes (Fig 2b-d). For energetically efficient transportomes, organisms therefore adopted different strategies, likely due to their specialisations and different requirements. The trend of expansion of ion channels and secondary transporters at the expense of ATP-dependent transporters also holds true for the prokaryotic transportomes (Fig 3), even though they did not undergo the kind of intensive developmental specialisation as was required in multicellular eukaryotes.

**Fig 3.**
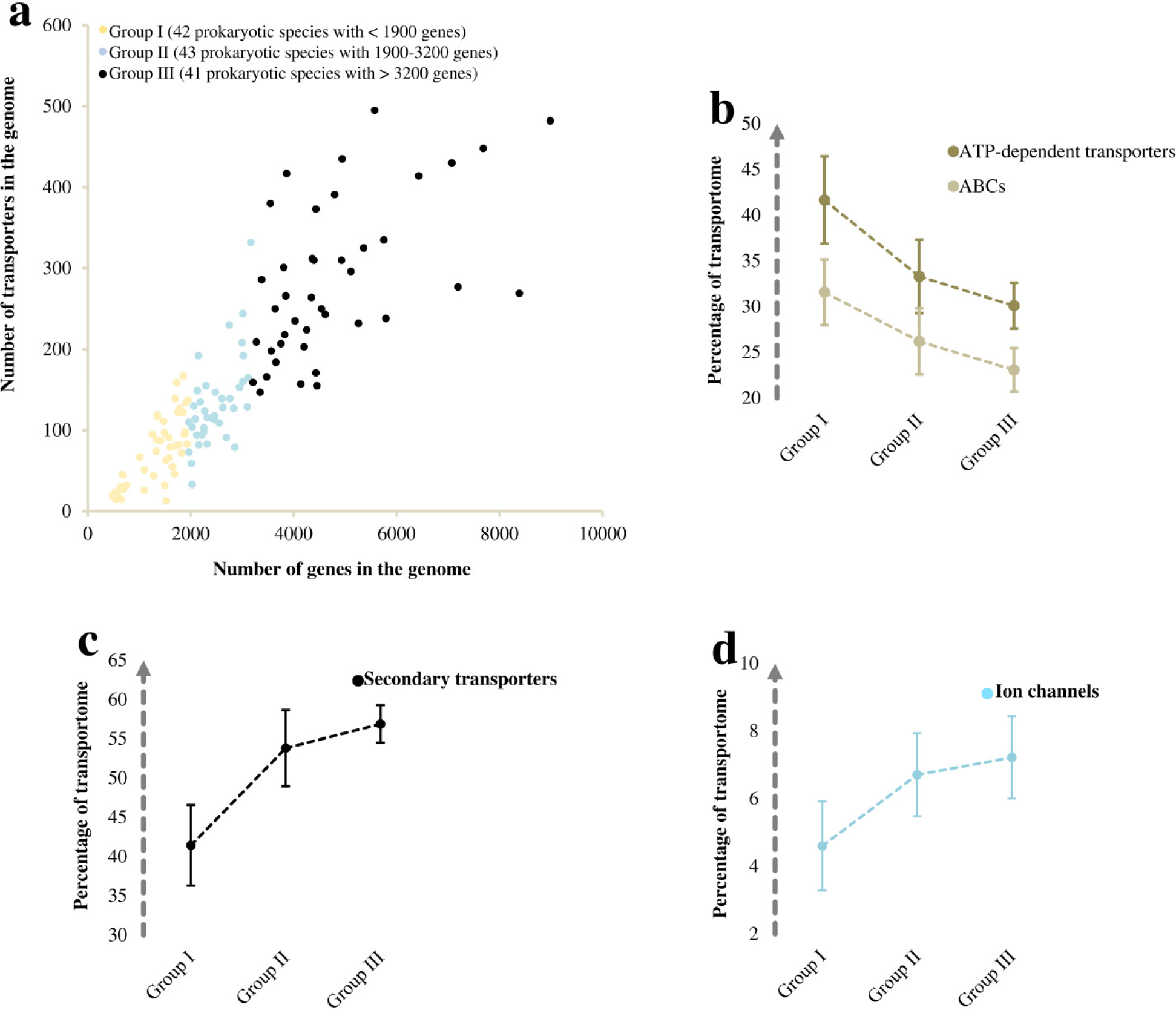
The compositional changes in the transportomes of prokaryotes with different genome sizes. **(a)** Comparisson between the transportome size and total gene number among prokaryotes including bacteria and archaea. All of the 126 studied species are clustered into three groups based on the total number of the genes. **(b)** The fraction of ATP-dependent transporters in the transportomes. **(c)** The fraction of secondary transporters in the transportomes. **(d)** The fraction of ion channels in the transportomes. The values on panels b-d are shown as mean +/-t-test based 99% confidence interval. The variations were also confirmed on the arc sin √*x* transformed data (See S1 Data). Group I and III differ significantly for all of the transporter classes with a *p*-value of < 0.001.

The analyses also indicate family member expansions for secondary transporters (from 60-100 in prokaryotes to more than 400 in animals and 600 in plants in average) and ion channels (from 10 members in prokaryotes to more than 300 members in animals). To find prokaryotic and eukaryotic specific families, we further enriched our analysis by including the publicly-available data for the secondary transporters and ion channels of prokaryotes found at the TransportDB 2.0 (http://www.membranetransport.org/transportDB2/index.html). This extended our prokaryotes to 2637 transportomes (S1 Table). We also included two eukaryotic transportomes from diatoms found at TransportDB 2.0 (S1 Table). When comparing the organisms for the presence of different transporter families, we found that eight secondary transporter families were completely lost in the passage to eukaryotes (See S1 Data). Additionally, we found that five new families of secondary transporters had emerged in eukaryotes (See S1 Data). Specifically, we did not find any prokaryotic hits in GenBank for these eukaryotic families, that must have diverged massively [26] from some ancestral genes. In contrary to secondary transporters, ion channels have evolved through both intra-family expansions and the substantial appearance of new families. While there were only seven families specific to the prokaryotes, 18 eukarya-specific families were found for ion channels (See S1 Data). Importantly, the mitochondria-specific solute carriers, *i.e.*, solute carrier family SLC25 [27,28], are absent from the transportomes of all 143 archaeal species (See S1 Table for the list of organisms). Among the bacteria, including 259 alpha-proteobacteria, of which 69 belonged to the order *Rickettsiales*, we found only seven bacterial genomes that encoded members of the mitochondrial transporter family. These are *Neorickettsia risticii*, *Neorickettsia sennetsu*, *Legionella pneumophila*, *Legionella longbeachae*, *Acidaminococcus intestini*, *Cardinium endosymbiont* and *Butyrivibrio proteoclasticus*. The presence of the mitochondrial transporter family members indicates that these seven bacterial species are possible origins of the mitochondrion in eukaryotes. The first two species are Gram-negative obligatory intracellular bacteria from the order *Rickettsiales*, an order that in previous studies was proposed (based on different evidence) as the most likely origin of the mitochondrion in eukaryotes [26–29].

A surprising insight here was related to the five evolutionarily younger secondary transporter families found only in eukaryotes (See S1 Data): three of these, about which we have experimental information, were energy-independent and low-energy-demanding transporter families for choline, silicate, and vitamin A. The last two belong to the 4 TMS multidrug endosomal transporter and chloroplast maltose exporter families. The choline transporter-like family is involved in choline influx [33]. The ‘birth’ of a cheap and sodium-independent transporter for choline is important because of the broad cellular usage of choline. Choline is an essential precursor for membranes and for the neuromodulator acetylcholine [34,35]. The silicon transporter family is also energetically cheap and has a silicate:sodium symport stoichiometry of 1:1 [36]. The animal-specific vitamin A receptor/transporter (STRA6) mediates costless influx and efflux of vitamin A derivatives by a mechanism not seen in any other transporter class [37,38]. Of particular interest, the animal visual system is dependent on vitamin A and has a substantial energetic cost, *e.g.*, up to 15% of resting metabolism in the Mexican fish *Astyanax mexicanus* [39,40]. Such photo-detection involves the single-photon-triggered isomerization of 11-*cis*-retinal to all-*trans*-retinal, which must be recycled back through efflux and influx steps of these isomers between the retinal pigmented epithelial and photoreceptor cells [37,41–43]. Thus, the evolved energy-independent membrane translocation of the vitamin A isomers seems to be an adaptive trait for higher energetic performance. This is in line with the positive selection reported for STRA6 in different mammalian phyla [44]. Additionally, we found a higher representation of ion channels in the animal kingdom when compared to the other domains of life (Fig 2d). The membrane transport of ions is important for highly energy-demanding sensory tissues [7,45] and it can therefore be hypothesized that the extensive diversification of ion channels and the costless transport of vitamin A in animals are trade-offs between the benefits of the evolved nervous and visual systems and their high energy requirements. In conclusion, our results provide genome-scale molecular evidence to the evolution of the cellular transportome towards an improved energetic efficiency.

## Materials and Methods

The publicly-available membrane transporter data on ion channels and secondary transporters were extracted from TransportDB (http://www.membranetransport.org/transportDB2/) [22]. The transportomes of 126 prokaryotic species (78 bacteria and 48 archaea) and 96 eukaryotic species (22 primitive eukaryotes, 24 algae and plants, 23 fungi, and 27 animals) (See S1 Data) were annotated using the Transporter Automatic Annotation Pipeline at TransportDB [22]. We also included 27 eukaryotic species including transportomes of 8 primitive eukaryotes, 6 algae and plants, 7 fungi, and 6 animals publicly available at TransportDB (See S1 Data). To study the compositional changes of transportomes, we did not include any transportomes from prokaryotes publicly available at the TransportDB. This is due to the incomplete information on the ABC transporters. The majority of ABC transporters in prokaryotes are coded by different genes of an operon, where each gene codes for different subunits [46], and these should be excluded from the data and considered as single transporter in our analyses on the transportome composition. However, this information on the ABC coding genes and operons is not provided for the publicly-available transportomes of prokaryotes.

To predict the transportomes, the proteomes of organisms were downloaded from the Genbank and Ensembl databases (http://www.ensembl.org/index.html, http://fungi.ensembl.org/index.html, http://protists.ensembl.org/index.html, http://plants.ensembl.org/index.html) [47]. All proteins with fewer than 100 amino acids were excluded. Taken together, we analysed 78 bacterial, 48 archaeal, 30 primitive eukaryotes, 30 algal and plant, 30 fungal, and 33 animal transportomes, each representing one independent biological replicate (per species). The list of organisms with their genome size and total number of transporters is shown in S1 Data and S1 Table. The annotated transportomes were manually filtered for multi-prediction hits as well as the alternative isoforms of transport proteins before the analysis. Taken together, our analyses on the transporter families included all of the 15 ATP-dependent transporter families, 51 ion channel families, and 90 secondary transporter families which were present in the studied organisms. We used Student’s t-test to examine the possible differences among the samples, *i.e.*, domains of life. The data were in the format of counts and percentages. Therefore, to exclude a possible divergence from normality, we also performed the analyses on the arc sin √x transformed data. All the analyses can be found in S1 Data.

To make an approximation of the energetic performance of the cellular transportomes, we calculated the inter-transportome changes in energy-usage efficiency (EUE), which we defined as the average free energy demand for a single transport event. The following assumptions were made when calculating EUE values. Equilibrative ion channels require no free energy for the transport action. While some of the secondary transporters also act in an energy-independent manner known as equilibrative transport or facilitated diffusion, others exploit the electrochemical potential established by ATPases across membranes [13,21,48–50]. For active transport, secondary transporters are considered to use membrane electrochemical potential gradients, coupled to a varying stoichiometry (0.5 to 3) of ions per turnover [49,51–72]. Considering the stoichiometry of ≈ 2-3 ions pumped per ATP hydrolysed by ATPases [73–82], even concentrative secondary transporters would not use more than 0.5 ATP per substrate translocation across a membrane on average. The average of two substrates per ATP tends, therefore, to be conservative for secondary transporters and a higher rate of substrate translocation can also be expected. In contrast, the ATP-dependent members belonging to the ABC superfamily, and also the mitochondrial protein translocase, the type III secretory pathway, the chloroplast envelope protein translocase, and the arsenite-antimonite efflux families, generally show a 1:2 stoichiometry of substrate:ATP hydrolysis [83–89]. To calculate the inter-transportome variations in the energy-usage efficiency (∆EUE), we therefore applied the equation: 
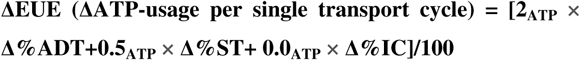
where ADT, ST, and IC are ATP-dependent transporters, secondary transporters, and ion-channels, respectively. The F/V/A-type ATPases and ATP synthases were excluded from the ATP-dependent transporters. This is because they provide the energy as ATP or membrane electrochemical potential for the rest of the transporters. It is worthy of note that the efficiency of energy usage indicates the average energy demand per unit of action. So the expression and activity levels of transporters are not taken into the account when calculating the EUE. A more general example would compare two organisms with exactly the same transportome but with differences in the expression levels of transportome members among them. Here, the EUE of transportome machineries of the two organisms are equal since they use exactly same transportome machinery. This is thus irrespective of the differences in activity levels of transporters, which define the total energy demand for the given transportome.

## Abbreviations

ADT: ATP-dependent transporters
EUE: energy-usage efficiency
IC: ion-channels
ST: secondary transporters

## Availability of data

The datasets supporting the conclusions of this article are included within the article and Supplementary Data 1.

## Supplementary Information

### Supplementary Table 1

The list of organisms with publicly-available data on the ion channels and secondary transporters.

### Supplementary Data 1

Genomic and transcriptome data on the organisms included in the study.

## Author contributions

BD, DBK, and IB conceived the study. BD performed the analyses by consulting DBK and IB. BD, DBK, and IB wrote the manuscript.

## Competing financial interests

The authors declare no competing financial interest.

## Acknowledgements

B.D. and I.B. acknowledge the financial support by the Novo Nordisk Foundation. D.B.K. thanks the Biotechnology and Biological Sciences Research Council (grants BB/M006891/1, BB/M017702/1 and BB/P009042/1) for financial support.

